# α-Parvin Promotes Glucose Uptake and Metabolism in Skeletal Muscle with Minimal Influence on Hepatic Insulin Sensitivity

**DOI:** 10.1101/2025.09.22.676818

**Authors:** Fabian Bock, David A. Cappel, Xinyu Dong, John W. Deaver, Dan S. Lark, Luciano Cozzani, Deanna P Bracy, Louise Lantier, Kakali Ghoshal, Allison Do, Richard L Printz, Owen P. McGuinness, David H. Wasserman, Ambra Pozzi, Roy Zent, Nathan C. Winn

## Abstract

Skeletal muscle and liver insulin resistance are early features in the sequelae of type 2 diabetes. Integrins are extracellular matrix receptors expressed on skeletal muscle cells and hepatocytes and have been implicated in modulating obesity-associated insulin resistance. Integrins regulate cell function through intracellular proteins including the ILK-PINCH-Parvin (IPP) complex. ILK promotes skeletal muscle and liver insulin resistance in diet-induced obesity in mice but the role of Parvin is unexplored. Here we demonstrate that hepatocyte specific deletion of α-Parvin had only minimal influence on endogenous glucose production or whole-body insulin sensitivity. In contrast, deletion of α-Parvin in skeletal muscle caused a striking reduction in muscle glucose uptake during an insulin clamp in lean mice which was not exacerbated by diet-induced obesity. Insulin-mediated GLUT4 membrane recruitment was impaired in mutant muscles which displayed significant morphological abnormalities due to actin cytoskeleton dysfunction. Consistent with severe muscular dysfunction, mitochondrial oxidative capacity and aerobic exercise capacity were blunted in muscle α-Parvin-null mice. Thus, α-Parvin has a minor role in liver insulin action but is required for insulin-stimulated glucose uptake in skeletal muscle due to its role in actin cytoskeleton regulation. These data suggest that individual IPP complex proteins link cell structure to metabolism via distinct mechanisms in a tissue-specific fashion.

## Introduction

The obesity epidemic has increased the prevalence of insulin resistance, Type 2 Diabetes, and the Metabolic Syndrome (1–4). Liver and skeletal muscle (SkM) metabolic dysfunction are early and prominent defects in the sequalae of these conditions (5–14). Following a carbohydrate-rich meal, approximately 60% of glucose clearance from the blood is attributable to the combined actions of liver and SkM in insulin sensitive humans (15). Impaired glucose regulation in SkM and liver is a central feature of obesity induced metabolic dysfunction. The molecular mechanisms that contribute to this loss of function are not fully defined, which may limit clinical treatment options.

Liver and SkM cells are in contact with extracellular matrix (ECM) components and have a defined morphology that is determined by the intracellular actin cytoskeleton (16, 17). Our group has shown that the family of adhesion receptors, known as integrins, are key regulators of metabolic function in the muscle (18–20). Integrins are heterodimeric transmembrane receptors comprised of α and β subunits. Integrins bind the extracellular matrix, coupling it to cytoskeletal dynamics and signal transduction by recruiting intracellular binding proteins to their cytoplasmic tail (21). The ILK-PINCH-Parvin (IPP) complex is a well characterized regulator of integrin function that links the cytoplasmic domains of integrins with cytoskeletal components (22, 23).

Deletion of ILK or PINCH results in the dissolution of the IPP complex (24, 25), suggesting that the stability of the complex is dependent on the presence of the members of the complex. The IPP protein Parvin has α, β, and LJ isoforms, of which α-Parvin is the primary SkM isoform (26). α-Parvin localizes to focal adhesions (FAs) where it interacts with filamentous actin (F-actin) through their actin-binding domain (27), as well as with actin-regulatory proteins including regulators of Rho GTPases (28, 29). The principal function of α-parvin is to regulate actin polymerization, which is mediated by the Rho GTPases and in many situations, deletion of α-parvin has phenocopied those observed when ILK was deleted.

Studies show that ILK is a driver of insulin resistance in SkM and the liver (19, 30–32). A key question is whether other IPP complex components have similar, overlapping, or different metabolic functions. In this study, we generated SkM-selective and hepatocyte-selective α-Parvin knockout mice to test the role of α-Parvin in SkM and liver insulin resistance. We show that α-Parvin deficiency impairs glucose uptake and oxidative metabolism in SkM with minimal influence over liver insulin action. These findings indicate that the mechanism by which α-Parvin regulates muscle metabolism is distinct from that seen in mice with ILK deletion (19). Thus, individual IPP complex proteins link cell structure to metabolism in a tissue-specific fashion via distinct mechanisms.

## Materials and Methods

All procedures were approved and conducted in compliance with the Vanderbilt University Institutional Animal Care and Use Committee. Vanderbilt University is accredited by the Association for Assessment and Accreditation of Laboratory Animal Care International.

### Mouse Models

Mice were housed in a temperature (∼22°C) and humidity-controlled facility with a 12-hour light cycle under supervision of the Vanderbilt University Division of Animal Care. To isolate the role of α-Parvin on hepatic and SkM insulin resistance, we developed hepatic (hParKO) and SkM (mParKO) specific α-Parvin knockdown mouse models. α-Parvin^f/f^ mice (generously provided by Reinhard Fässler) were crossed with mice expressing *cre* recombinase under control of the albumin promoter (Strain #:003574, Jackson Laboratory (33)) to target hepatocytes or SkM specific human skeletal actin promoter (Strain#:006149, Jackson Laboratory (34)) to target myocytes. All mice were on the C57BL/6J strain. Par^f/f^ mice were used as controls. At 6 weeks of age, male mice were fed standard chow diet (5001 Laboratory Rodent; LabDiet) or 60% fat diet (HFD, Research Diets, #D12492, 5.21 kcal/g food) for 12 weeks with experiments performed between 16 and 18 weeks of age.

### Body composition

Mouse body fat mass, fat-free mass (FFM), and free water were measured by a nuclear magnetic resonance whole-body composition analyzer (Bruker Minispec).

### Hyperinsulinemic-Euglycemic Clamp

Catheters were surgically placed in a carotid artery and jugular vein for sampling and infusions, respectively, one week before clamp procedures (35). Mice were transferred to a 1.5 L plastic container without access to food 5 hour prior to the start of an experiment. Two independent euglycemic clamp studies were conducted for the Liver and SkM muscle experiments, respectively.

*Study 1 (Liver Parvin Experiments):* At 7 am (t=-300), catheters were attached to extensions and secured to syringes. ^2^H_2_O (2.7 µl/g body weight) was given to enrich body water to 4.5%.

All infusates were given in salinized 4.5% ^2^H_2_O to prevent dilution of the body ^2^H_2_O pool. After 3 hours of fasting, an arterial blood sample was obtained to determine natural isotopic enrichment of plasma glucose. At 90 min prior to initiation of the clamp, a quantitative stable isotope delivery to increase glucose isotopic enrichment above natural isotopic labelling was initiated. [6,6-^2^H_2_]glucose was primed (16 mg) and continuously infused for a 90 min equilibration and basal sampling periods (0.8 mg/kg/min in saline). The insulin clamp was initiated at t=0 min with a continuous insulin infusion (2.5 or 4 mU/kg body weight/min for lean and diet-induced obesity (DIO) mice, respectively), which continued for 145 min. Arterial glucose was clamped using a variable infusion rate of glucose + [6,6-^2^H_2_]glucose (0.08 mass percent excess), which was adjusted based on the measurement of blood glucose at 10 min intervals during the 145 min clamp period. By mixing the glucose tracer with the unlabeled glucose infused during a clamp, deviations in arterial glucose enrichment are minimized and steady state conditions are achieved. Mice received heparinized saline-washed erythrocytes from donors at 5 μl/min to prevent a fall in hematocrit. Baseline blood or plasma variables were calculated as the mean of values obtained in blood samples collected at −15 and −5 min. Blood was taken from 80– 120 min for the determination of plasma enrichment. Clamp insulin was determined at t=100 and 120 min. After the last sample, mice were anesthetized and tissues were freeze-clamped for further analysis.

Study 2 (Muscle Parvin experiments): [3-^3^H]glucose was primed and continuously infused from t=-90min to t=0 min (0.06 µCi·min^-1^). The insulin clamp was initiated at t=0 min with a continuous insulin infusion (4 mU·kg^-1^·min^-1^) and variable glucose infusion initiated and maintained until t=155 min. The glucose infusate contained [3-^3^H]glucose (0.06 µCi·µl^-1^) to minimize changes in plasma [3-^3^H]glucose specific activity. Arterial glucose was monitored every 10 min to provide feedback to adjust the glucose infusion rate as needed to maintain euglycemia. Erythrocytes were infused at a rate calculated to compensate for blood withdrawal over the duration of the experiment. [3-^3^H]glucose specific activity was determined at -15 min and -5 min for the basal period, and every 10 min between 80 to 120 min for the clamp period to assess glucose disappearance and endogenous glucose production. At 120 min, A 13 µCi intravenous bolus of 2-[^14^C]-deoxyglucose ([^14^C]2DG) was administered at 120 min to determine the tissue glucose metabolic index (Rg), an index of tissue-specific glucose uptake (36–38).

Blood samples were collected at 122, 125, 130, 135 and 145 min to measure plasma [^14^C]2-DG. [^14^C]2DG is phosphorylated in tissue by a hexokinase but is not a substrate for further metabolism. The accumulation of the isotopic phosphorylated glucose analog, [^14^C]2DGP, in excised tissue is used to calculate tissue Rg.and blood At 145 min, mice were euthanized and tissues immediately harvested and freeze-clamped.

### Plasma and Tissue Sample Processing and Glucose Flux Rate Determination

Plasma glucose concentrations were measured using commercial colorimetric assay (Sigma). Plasma insulin was measured by radioimmunoassay (Sigma-Aldrich) by the MMPC Analytical Resources Core. Plasma was derivatized to obtain di-O-isopropylidene propionate derivative of glucose, and plasma glucose enrichments ([6,6-^2^H_2_]glucose) were assessed by GC-MS, as described previously (39). Glucose rate of endogenous appearance and rate of disappearance rates were determined using steady-state equations (40). Endogenous glucose appearance was determined by subtracting the glucose infusion rate from total rate of appearance.

Gluconeogenesis and glycogenolysis were measured using positional glucose labeling from ^2^H_2_O, as described previously (41). The glucose metabolic index (Rg) was calculated as previously described (42). Plasma and tissue (gastrocnemius, vastus lateralis, gonadal adipose tissue, subcutaneous adipose tissue, brain, and heart) radioactivity for [3-^3^H]glucose, [^14^C]2DG and, [^14^C]2DG-6-phosphate was measured by scintillation counting of deproteinized samples as previously described. Radioactivity in plasma samples, and in tissue samples were determined by liquid scintillation counting. Glycogen assay was performed according to Chan and Exton (43).

### Quantification of Liver Lipids

Liver lipids were extracted using the method of Folch-Lees (44). The extracts were filtered and lipids recovered for separation by thin layer chromatography as previously described in the chloroform phase. Individual lipid classes were separated by thin layer chromatography using Silica Gel 60 A plates developed in petroleum ether, ethyl ether, acetic acid (80:20:1) and visualized by rhodamine 6G. Phospholipids, diglycerides, and triglycerides were scraped from the plates and methylated using BF3 /methanol as described by Morrison and Smith (45). Gas chromatographic analyses were performed on an Agilent 7890A gas chromatograph equipped with flame ionization detectors and a capillary column (SP2380, 0.25 mm x 30 m, 0.20 µm film, Supelco, Bellefonte, PA). Helium was used as the carrier gas. The oven temperature was programmed from 160 °C to 230 °C at 4 °C/min. Fatty acid methyl esters were identified by comparing the retention times to those of known standards.

### Indirect calorimetry and ambulatory activity

Respiratory gases, locomotor activity, and feeding were determined using the Promethion Metabolic Analyzer (Sable Systems, North Las Vegas, NV). Rates of energy expenditure were calculated from VϑO_2_ and VϑCO_2_ using the Weir equation [EE k cl J^r1^) = 60 · (0.003941 ·v a_2_) + (0.001106 · v ca_2_]]. Mice were placed in metabolic cages, singly housed, for a total of 14 days. Day 0-6 data collection occurred in the presence of a locked running wheel. On day 7, the wheel was unlocked to determine voluntary running activity and changes in additional metabolic parameters (e.g., EE, food intake, non-exercise movement) when mice are allowed to be physically active.

### Incremental Exercise Stress Test

Stress tests were conducted on a single-lane treadmill from Columbus Instruments beginning at a speed of 10 m·min^-1^. Speed was increased by 4 m·min^-1^ every three minutes until exhaustion (46).

### High-Resolution Respirometry in Permeabilized Muscle Fibers

Oxidative metabolism was assessed by measuring oxygen consumption in *ex vivo* muscle fiber bundles isolated from the gastrocnemius muscle using an Oroboros Oxygraph2K system, as described (47). This method retains the functional interaction of mitochondria with normal cell architecture (48), which is lost in isolated mitochondria (49, 50). Muscle samples were collected from mice and held in a BIOPS solution (2.7 mM EGTA calcium buffer, 20 mM imidazole, 20 mM taurine, 50 mM K-MES, 6.56 mM MgCl_2_, 5.77 mM Na_2_ATP, and 15 mM phosphocreatine, pH 7.1). Muscle fiber bundles weighing approximately 5-10 mg were dissected away from each other and surrounding connective tissue using sharpened forceps and then permeabilized with a treatment of 50 μg/mL saponin in BIOPS for 30 minutes. Tissues were washed for a minimum of 15 minutes in Mir05 Solution (30mM KCl, 105mM K-MES, 10 mM K_2_HPO_4_, 5mM MgCl_2_-6H_2_O, 1mM EGTA, 0.5g/L BSA). After the wash bundles were placed in the respirometer wells containing Mir05 solution. Wells were hyperoxygenated to 250 nmol/mL by injection of O_2_ into the wells. Oxygen consumption was normalized to the wet weight of the fiber bundles. Three different combinations of substrates were used to measure complex 1 activity (10mM glutamate, 2mM Malate, 2mM ADP), complex 1&2 activity (10mM glutamate, 2mM Malate 10mM Succinate, 2mM ADP), and oxygen consumption derived from fatty acid oxidation (2mM Malate, 2mM ADP, 75µM Palmitoylcarnitine).

### Cell Imaging and Immunohistochemistry

Liver specimens were fixed in 10% formalin paraformaldehyde in phosphate buffered saline and embedded in paraffin. Paraffin tissue sections were stained with hematoxylin-eosin (H&E, Epredia, 7211 & 22-110-637) for evaluation of tissue structure. A group of mice were fasted for 5 hours followed by an i.p. injection of insulin (1U/kg body weight) to determine the occupancy of GLUT4 in the muscle membrane. For this study mice were euthanized 10 minutes post insulin injection. Sections of gastrocnemius muscle were collected from mice and fixed in a solution of 10% formalin paraformaldehyde in phosphate buffered saline or snap frozen in isopentane at liquid nitrogen temperature and affixed to cork blocks with OCT. Fixed tissues were embedded in paraffin and sectioned and frozen tissues were cryo-sectioned and mounted at the Translational Pathology Shared Resource at Vanderbilt University. Antibodies against Cav3 (Santa Cruz #sc55518), GLUT4 (Abcam #ab33780), α-Parvin (Cell Signaling #4026), were used and F-actin was visualized using AF-647–phalloidin (Thermo, #A22287). Additional slides were stained with H&E to visualize tissues. Muscle central nuclei percentage was determined by counting of cells with and without central nuclei throughout a whole slide. Slides were scanned at the Digital Histology Shared Resource at Vanderbilt University using a high throughput Leica SCN400 Slide Scanner automated digital image system from Leica Microsystems or imaged at the Cell Imaging Shared Resource. Confocal muscle images were collected with confocal microscopy at super-resolution using a Zeiss LSM 980 confocal microscope equipped with an inverted Axio Observer 7 and Airyscan 2 detector. The objective used was a 63×/1.4 numerical aperture (NA) Plan Apochromat oil or 10×/0.50 NA Plan Apochromat (for low-powered scanning of F-actin-labeled muscle fibers). Airyscan super-resolution images were acquired under identical settings for all groups and images. Acquisition and 2D Airyscan processing of acquired images was done using ZEN Blue software (Carl Zeiss). Line Scan profiles of fluorescence intensity along an annotated line was performed using the plot profile function in Fiji/ImageJ.

### Immunoblotting

Immunoblotting was performed on basal and insulin-stimulated muscles. Mice were fasted for 5 hours followed by an i.p. injection of insulin (1U/kg body weight) or saline. After 10 minutes, mice were euthanized and muscle tissue was snap frozen for western blotting. Protein levels for Cofilin, p-Cofilin, Akt, p-AKT(Ser473), p-AKT(Thr308), and HKII were determined by immunoblotting using antibodies from Cell Signaling Technologies. Whole-tissue lysates were obtained by homogenization in lysis buffer. Mitochondrial and cytosolic fractions were obtained, as previously detailed (51). 30 ug of protein extract from gastrocnemius muscle were loaded onto Criterion TGX stain-free precast gels and transferred to nitrocellulose membrane using the Trans-Blot Turbo Transfer System (Bio-Rad). Total protein was measured using the stain-free ChemiDoc imaging system (Bio-Rad) and quantified using Image Lab software (Bio-Rad).

Membranes were blocked with Intercept blocking buffer (Licor) and then incubated with primary antibodies diluted 1:1000 in intercept blocking buffer. Anti-mouse or Anti-rabbit secondary antibodies conjugated to either a 700 or 800 nm florescent probe (Licor) diluted 1:10000 in intercept blocking buffer were used to visualize proteins on an Odyssey scanner at the Vanderbilt Molecular Biology Core Facility. Protein levels on blots were quantified by densitometry using Image Lab software (BioRad).

### Rho GTPase and Rac1 Activity Assay

Activation of Rho GTPases was measured using a commercially available ELISA assay (Cytoskeleton Inc). Antibodies for the active forms of RhoA and Rac1 were used to detect the levels of active rho GTPases. The amount of active protein was determined using a HRP-conjugated secondary antibody that was detected by absorbance at 490 nM.

### Statistical Analysis

T-tests were run to determine differences between Par^f/f^ and ParKO mice. If data did not follow a Gaussian distribution, non-parametric Mann-Whitney tests were used to determine statistical significance. In experiments that contained more than two groups, one-way analysis of variance (ANOVA) or two-way ANOVA models were conducted with pairwise comparisons using Tukey or Sidak correction. Brown-Forsythe correction was applied to groups with unequal variance.

For metabolic cage data, a generalized linear model with analysis of covariance was run to determine statistically significant differences via CalR online resource (https://calrapp.org/<u>#</u>). A linear regression model was used to test differences in cumulative food intake and wheel running distance. Data are presented as mean ± standard error (SE). An adjusted p value of <0.05 was used to determine significance.

## Results

### Liver deletion of ***α***-Parvin has mild effects on insulin action

To test the role of liver alpha-Parvin in diet-induced obese (DIO), we generated liver specific Parvin KO mice by crossing Par^f/f^ with Albumin-Cre mice (Fig 1A). Par^f/f^ and hParKO mice were placed on chow orϑ HFD for 12 weeks starting at 6 weeks of age to generate lean and DIO conditions (Fig. 1A). α-Parvin protein expression was markedly decreased in liver from hParKO mice compared to Par^f/f^ controls (Fig. 1B). This reduction was also associated with a 35% and 15% decrease in ILK and PINCH, respectively in hParKO livers (Supplemental Fig. 1). No differences in body weight were found between genotypes in lean or DIO conditions (Fig. 1C). We evaluated liver tissue structure by H&E staining and found that steatosis was markedly increased in DIO versus lean mice. The lipid accumulation was primarily in the form of microvesicular steatosis and did not appear to be different between hParKO and Par^f/f^ mice (Fig. 1D). To confirm this visual observation, we quantified total triglycerides, diglycerides, and phospholipids in lean and DIO Par^f/f^ versus hParKO mice. DIO increased liver triglycerides and diglycerides, whereas phospholipids were decreased. The lack of hepatic α-Parvin did not improve or worsen the liver lipid content in lean or DIO groups (Fig. 1E).

**Figure 1.**
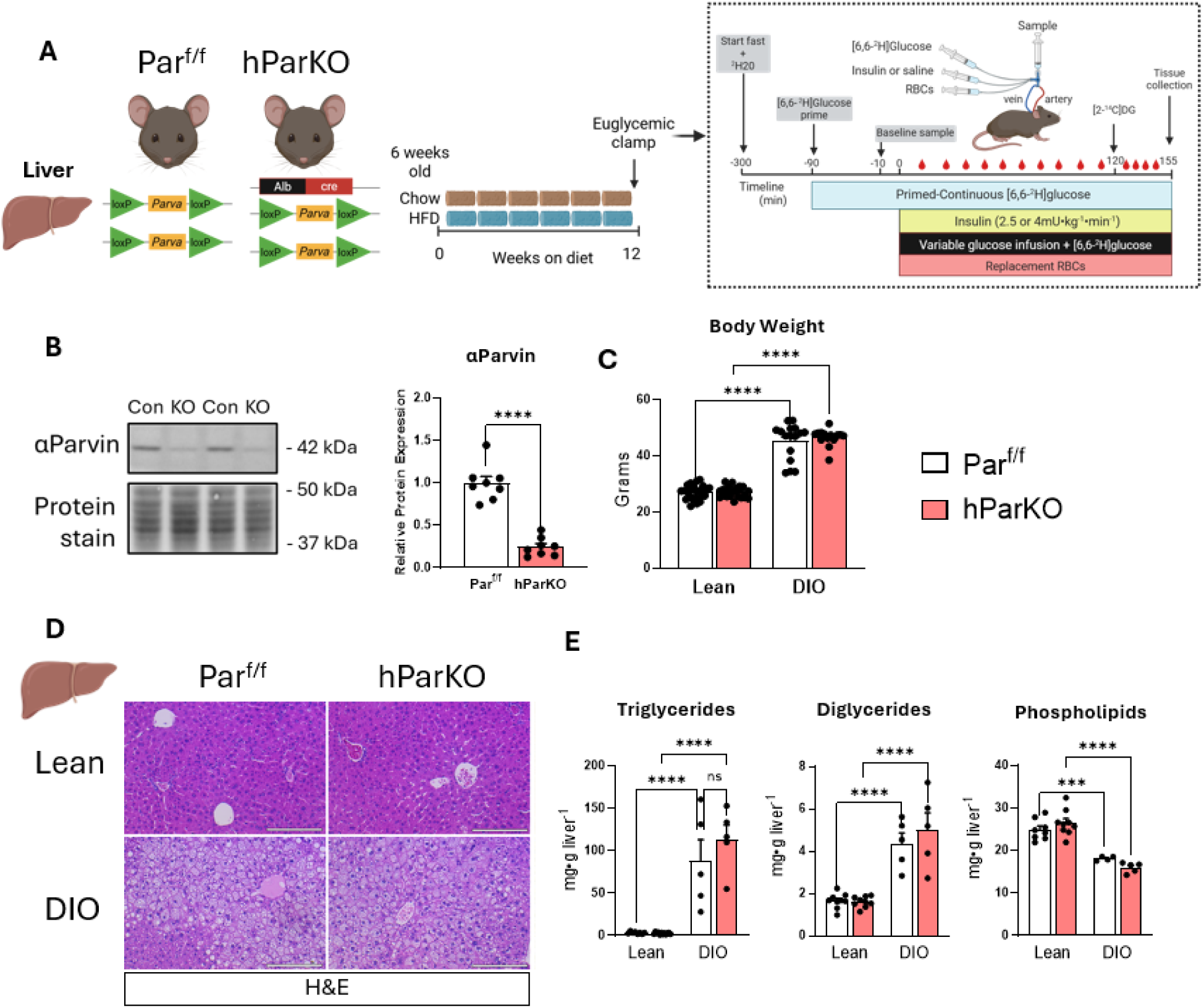
–. Hepatic _α_Parvin has minimal influence on liver morphology and steatosis in lean and DIO mice. **A**) Par^f/f^ mice were crossed with Albumin-Cre mice to generate deletion of _α_Parvin in hepatocyte (hParKO). At 6 weeks of age mice were fed chow or HFD for 12 weeks to generate lean and DIO groups. Mice underwent an hyperinsulinemic-euglycemic clamp after 12 weeks of diet intervention. **B**) immunoblotting of _α_Parvin in liver lysates was performed. Parvin expression was normalized to total protein using stain free gels containing covalent protein compound trihalo. **C**) Terminal body weight after 12 weeks of diet feeding. **D)** Liver sections were stained with H&E for evaluation of liver morphology. **E**) Livers were excised and snap frozen after a 5 hour fast. Liver lipids were extracted using the FOLCH method. Individual lipid classes were separated by thin layer chromatography and quantitative analysis performed using gas chromatography. Data are presented as mean ± SE. n=5-27 mice/group. Panel B, T test were conducted to test differences. Panel C and E, Two-way ANOVA with diet and genotype a factors was run to test for statistical differences. Panels D,E,I,J, Two-way repeated measures ANOVA with time and genotype as factors were run. Panels F-H and K-M, Two-way ANOVA with insulin and genotype as factors were run to test for differences. Alpha was p<0.05. ***p<0.001, ****p<0.0001, ns, not significantly different.

To determine liver and systemic insulin action, the hyperinsulinemic-euglycemic clamp technique was performed. ^2^H_2_O coupled with [6,6-^2^H_2_]glucose were used as tracers to determine glucose turnover and the rate of glycogenolysis and gluconeogenesis [(GNG), Methods]. In lean mice, plasma glucose was maintained at euglycemia (∼130 mg/dl) during clamped insulin infusion (2.5 mU/kg body weight/min) (Fig. 2A). Euglycemia was achieved by the exogenous infusion of glucose, which was not different between Par^f/f^ and hParKO mice (Fig. 2B). The rate of endogenous glucose production from glycogenolysis or GNG was not different between hParKO and Par^f/f^ mice in the fasted state or during the insulin clamp (Fig. 2C). Glucose disappearance was increased during the insulin clamp but no differences were found between Par^f/f^ and hParKO during basal or insulin clamped states, demonstrating that systemic insulin action is not influenced by liver α-Parvin (Fig. 2D). Plasma insulin levels were similar between genotypes at baseline and during experimentally clamped hyperinsulinemia (Fig. 2E). Thus, loss of liver α-Parvin has no discernable influence on hepatic or peripheral insulin action in lean and otherwise healthy mice.

**Figure 2.**
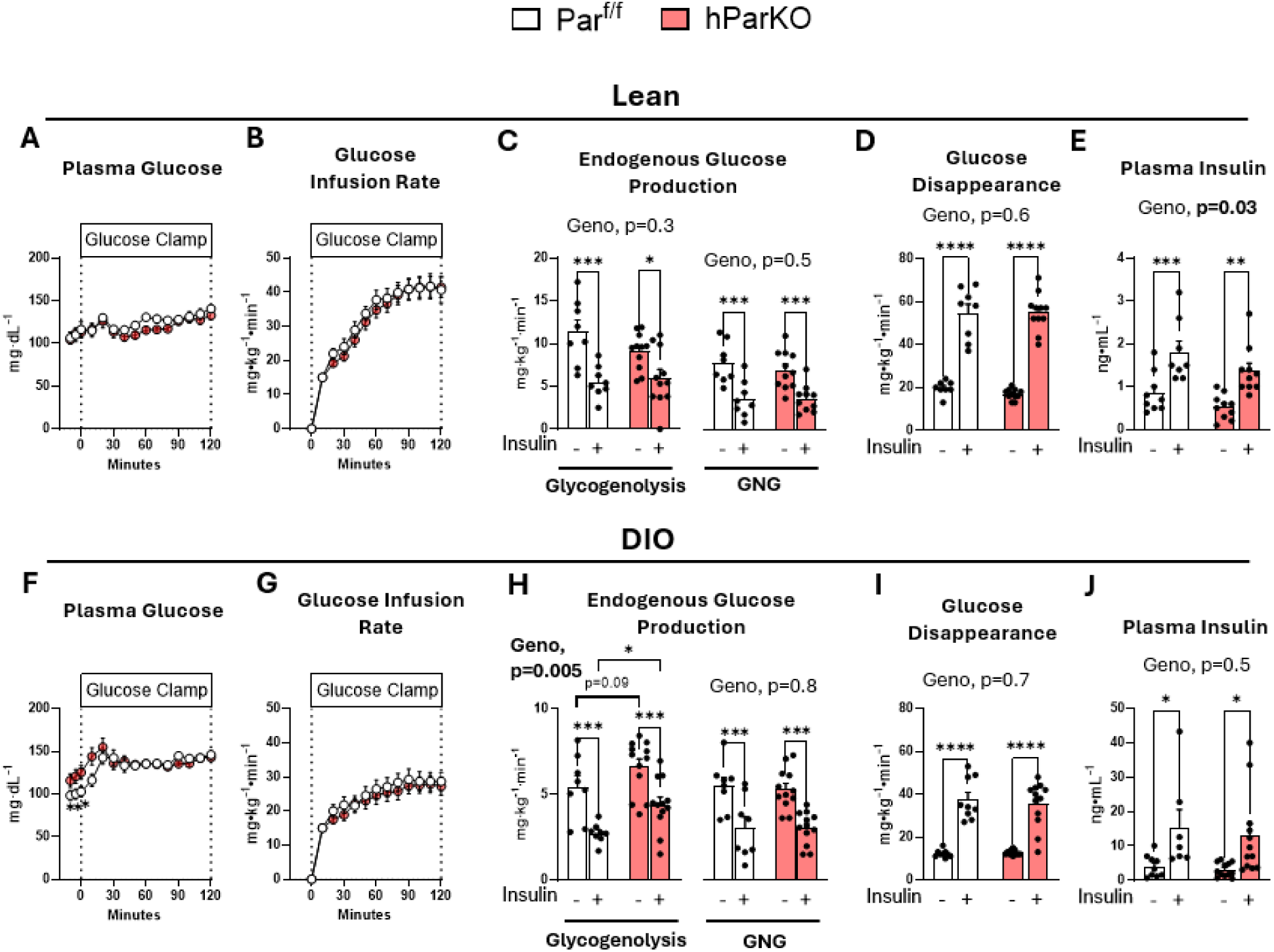
–. Hepatic _α_Parvin exerts a minor role on liver insulin action and glucose metabolism in lean and DIO mice. The hyperinsulinemic-euglycemic clamp technique was performed after 12 weeks of diet intervention in Lean (A-E) and DIO (F-J). At time=0 min a constant infusion of exogenous insulin was infused at 2.5 mU/kg/min and 4.0 mU/kg/min for lean and DIO groups, respectively. **A, F**) Plasma glucose levels were maintained at approximately 130 mg/dL during the glucose clamp by a **B, G**) variable glucose infusion rate to achieve steady state glucose levels. **C, H**) Endogenous glucose production rates were determined from [6,6-^2^H_2_] glucose using isotope dilution methodology. Infusion of ^2^H_2_O was performed to resolve rates of glycogenolysis and gluconeogenesis (GNG) from positional labelling of ^2^H. **D, I**) The glucose disappearance rate was determined from [6,6-^2^H_2_]glucose dilution. **E, J**) Plasma insulin levels were measured during fasting and steady state clamp conditions. Data are presented as mean ± SE. n=9-12 mice/group. Panels A,B,F,G, Two-way repeated measures ANOVA with time and genotype as factors were run. Panels C-E and H-J, Two-way ANOVA with insulin and genotype as factors were run to test for differences. Alpha was p<0.05. *p<0.05, **p<0.01, ****p<0.0001.

In DIO conditions, fasting glucose was ∼20% higher in hParKO vs Par^f/f^ mice (Fig. 2F). Plasma glucose levels were maintained at ∼130 mg/dL throughout the clamped insulin infusion (4 mU/kg body weight/min) (Fig. 2F). The glucose infusion rate required to achieve euglycemia was not different between DIO controls and hParKO mice (Fig. 2G). In the fasted state, the rate of endogenous glucose production from glycogenolysis was 20% higher in hParKO versus Par^f/f^ mice (Fig. 2H). Glycogenolysis was suppressed by insulin in both groups but remained significantly higher in hParKO mice (Fig. 2H). No significant differences in basal or insulin-stimulated rates of GNG were found and similarly glucose disappearance rates between genotypes were equivalent between the DIO Par^f/f^ and hParKO mice (Fig. 2H,I). Insulin levels during fasting and the clamp period were also not different between the genotypes (Fig. 2J). These data show that loss of hepatic α-Parvin in DIO mice elevates fasting glucose, due to the increased rate of glycogenolysis, with no difference in the response to insulin. In summary, hepatic insulin action is mildly affected by the loss of liver α-Parvin and this occurs in DIO mice.

### SkM deletion of ***α***-Parvin causes insulin resistance

We next determined whether deletion of α-Parvin in the SkM influenced insulin action (Fig. 3A). Par^f/f^ and mParKO mice were fed chow or HFD for 12 weeks at which time hyperinsulinemic-euglycemic clamps were performed (Fig. 3A). Immunofluorescence imaging of the gastrocnemius muscle collectively showed a ∼50% reduction in α-Parvin levels (Fig. 3B). ILK and PINCH levels, however, were not affected by the loss of α-Parvin (Supplemental Fig. 2). We found no difference in body weight in the mParKO mice versus Par^f/f^ mice on either chow or high-fat diet (Fig. 3C). An intraperitoneal glucose tolerance test was performed in lean and DIO mice to assess whether glucose homeostasis depends on muscle αParvin. We found a significant decrease in the rate of glucose clearance in lean mParKO mice versus lean controls (Fig. 3D). In contrast, no differences in glucose tolerance were found in the DIO Par^f/f^ versus mParKO mice (Fig. 3E). To determine the role of SkM α-Parvin in insulin sensitivity and glucose fluxes, hyperinsulinemic-euglycemic clamps were performed (Fig. 4). In both the lean and DIO mice the fasting blood glucose levels were similar between the mParKO and Par^f/f^ mice and were matched throughout the experiment (Fig 4A, F). The glucose infusion rate was slightly reduced in the mParKO mice in the chow but not in the HFD-fed condition (Fig 4B, G), indicating that insulin resistance is not amplified in mParKO mice when fed HFD. In lean mice, the endogenous rate of glucose production was not different between the genotypes, whereas glucose disappearance was decreased in response to insulin in mParKO mice, consistent with decreased peripheral insulin sensitivity (Fig. 4D). This reduction in insulin action was not related to changes in insulin levels at basal or during the clamp period (Fig. 4E). DIO mParKO mice were resistant to insulin-induced suppression of endogenous glucose production without showing differences in glucose disappearance compared with DIO controls (Fig 4H, I). Basal and clamp insulin levels were markedly elevated in DIO mParKO mice indicative of an insulin-resistant phenotype (Fig. 4J).

**Figure 3.**
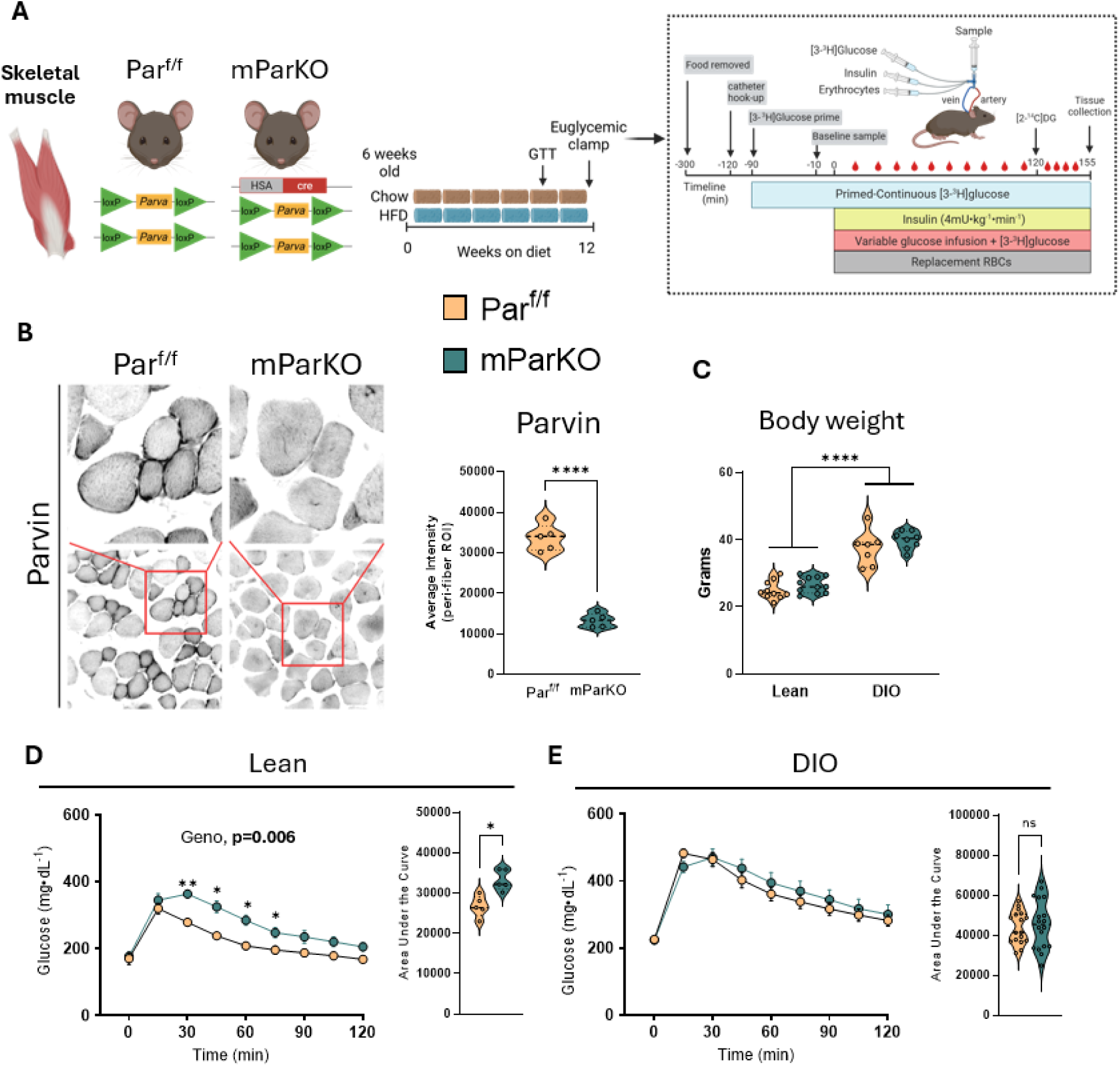
–. Skeletal muscle _α_Parvin improves glucose tolerance in lean mice. **A**) Parf/f mice were crossed with human alpha actin-Cre mice to generate deletion of _α_Parvin in skeletal muscle (mParKO). At 6 weeks of age mice were fed chow or HFD for 12 weeks to generate lean and DIO groups. The hyperinsulinemic-euglycemic clamp technique was performed after 12 weeks of diet intervention. **B**) immunofluorescence imaging of _α_Parvin was performed on OCT frozen sections of gastrocnemius muscle. The relative intensity was determined using ImageJ software. **C**) Terminal body weight after 12 weeks of diet feeding. Intraperitoneal glucose tolerance test (2g/kg body weight) was performed in **D**) lean and **E**) DIO mice after 10 weeks of diet feeding. Area under the curve was calculated and presented to the right of each glucose curve. Data are presented as mean ± SE. n=5-18 mice/group. Panel B and AUC from Panel D and E, T test were conducted to test differences. Panel C, Two-way ANOVA with diet and genotype as factors was run to test for statistical differences. Glucose curves in panel D and E, Two-way repeated measures ANOVA with time and genotype as factors were run. Alpha was p<0.05. *p<0.05, **p<0.01, ****p<0.0001.

**Figure 4.**
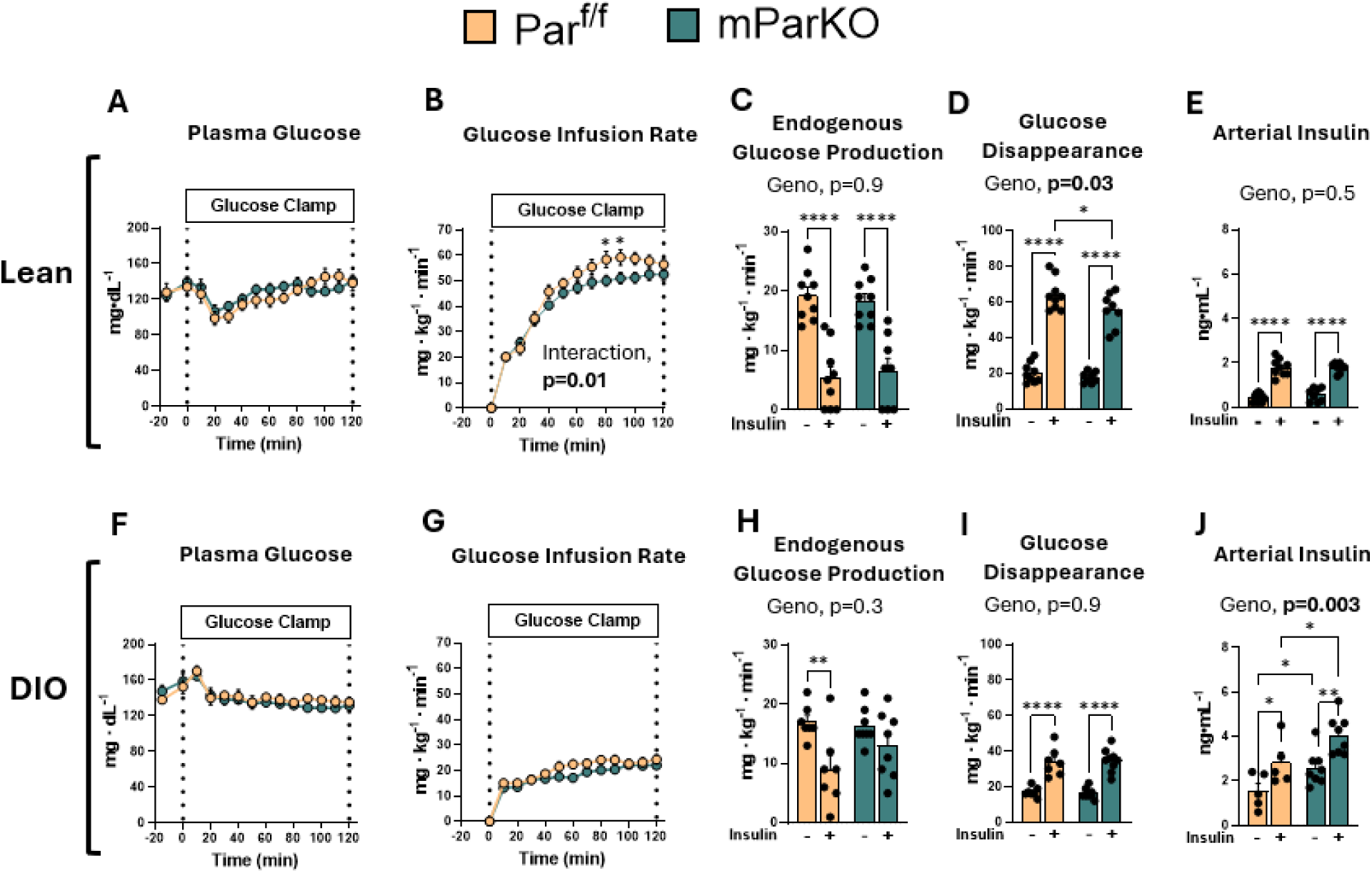
– Skeletal muscle _α_Parvin is protective against insulin resistance. The hyperinsulinemic-euglycemic clamp technique was performed after 12 weeks of diet intervention in Lean (A-E) and DIO (F-J). At time=0 min a constant infusion of exogenous insulin was infused at 4.0 mU/kg/min for both lean and DIO groups. **A, F**) Plasma glucose levels were maintained at approximately 130 mg/dL during the glucose clamp by a **B, G**) variable glucose infusion rate to achieve steady state glucose levels during hyperinsulinemia. **C, H**) Glucose rate of endogenous production and **D, I**) rate of disappearance were determined from [3-^3^H] glucose using steady-state equations. **E, J**) Plasma insulin levels were measured during fasting and steady state clamp conditions. Data are presented as mean ± SE. n=8-12 mice/group. Panel A-B, F-G, Two-way repeated measures ANOVA with time and genotype as factors were run. Panel C-E and H-J, Two-way ANOVA with insulin and genotype as factors was run to test for statistical differences. Alpha was p<0.05. *p<0.05, **p<0.01, ****p<0.0001.

We hypothesized that the decrease in glucose disappearance rate observed in lean mParKO mice (Fig. 4) would manifest as decreased tissue-specific glucose uptake (measured as the glucose metabolic index, Rg). Indeed, chow-fed mParKO mice showed ∼50% decrease in glucose Rg in the gastrocnemius and superficial vastus lateralis with no differences in adipose tissue, heart, or brain (Fig 5A). As expected, DIO worsened the insulin-stimulated glucose uptake in the muscle for both groups but did not exacerbate the metabolic phenotype of SkM deletion of αParvin (Fig 5B). The decrease in insulin-stimulated SkM glucose uptake was unrelated to Akt signaling, whereby the insulin-stimulated phosphorylation of Ser473 and Thr308 increased similarly in muscles of Par^f/f^ and mParKO mice (Fig. 5C). Hexokinase (HK) phosphorylates glucose thereby trapping it inside muscle cells. This represents the terminal step in SkM glucose uptake. Given the decrease in muscle Rg, which is directly dependent on HK activity, we determined whether HK expression was affected by the loss of α-Parvin. HK type II (HKII) constitutes the majority of total HK activity in SkM and is impaired in insulin resistance (52). HKII can dynamically move between the cytosol and outer mitochondrial membrane when growth factors, like insulin, initiate downstream signaling (53). This is significant because it is postulated that mitochondrial bound HKII funnels glucose derived carbon into the mitochondria coupling glycolysis to oxidative phosphorylation. Cytosolic and mitochondrial fractionation of muscles were prepared in basal and insulin-stimulated conditions, to determine whether loss of α-Parvin reduces the fraction of mitochondrial to cytosolic HKII. GAPDH and VDAC1 were used as indicators of cytosolic and mitochondrial cellular fractions, respectively. No differences in cytosolic HKII expression were found between Par^f/f^ and mParKO mice (Fig. 5C). In contrast, mParKO mice had reduced expression of mitochondrial-associated HKII (Fig. 5C), suggesting that dysregulated compartmentation of HKII in mParKO mice may influence carbon flux at the mitochondrial membrane and/or amplify deficits in glucose uptake. (Fig. 5D). The decrease in HKII in the mitochondrial fraction was associated with elevated SkM glycogen in mParKO mice (Fig. 5E). Collectively, these data reveal that SkM αParvin opposes insulin resistance.

**Figure 5.**
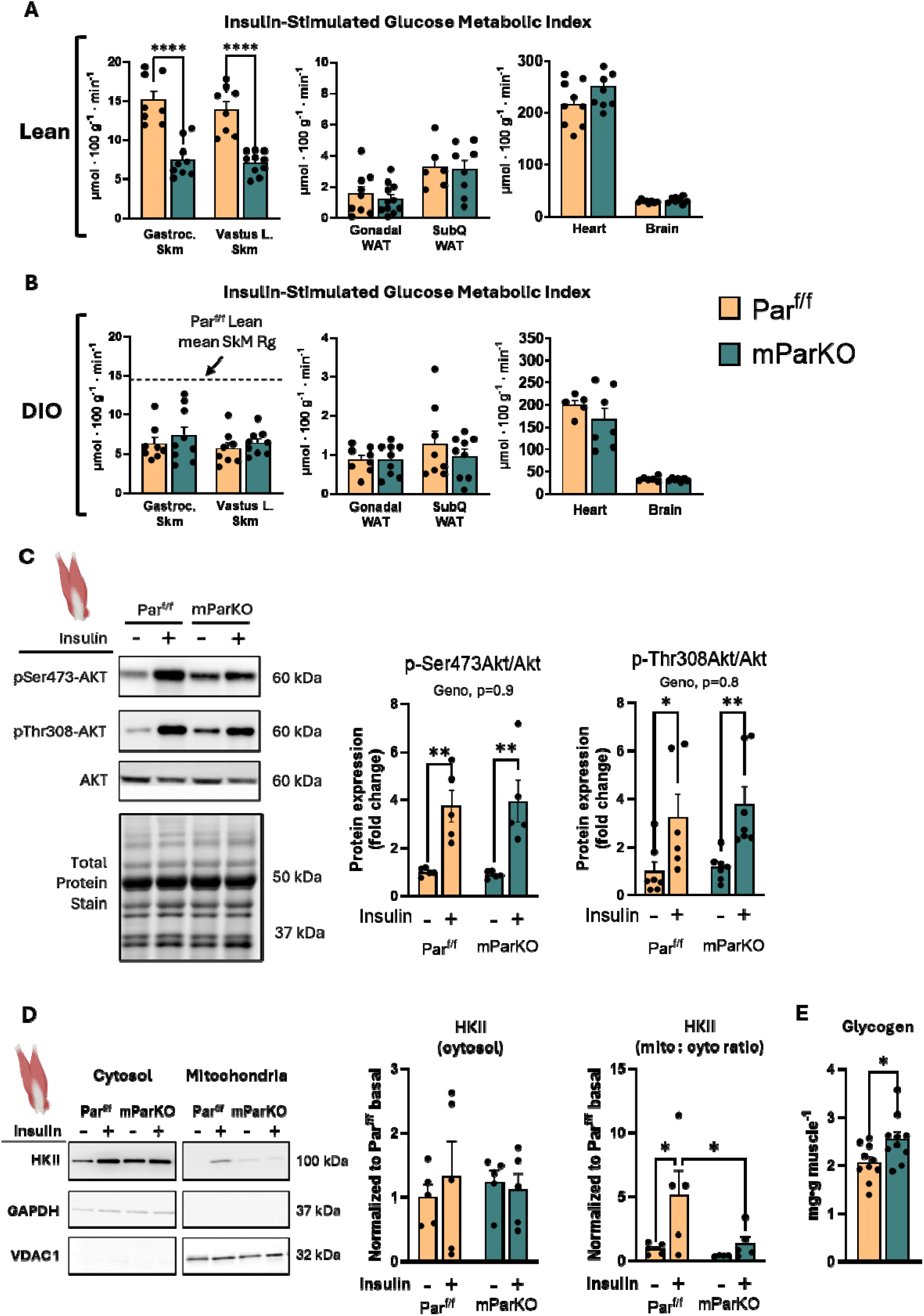
– Skeletal Muscle _α_-Parvin regulates muscle glucose uptake in lean mice. A bolus of 2[^14^C]-deoxyglucose was delivered during the steady state insulin clamp to determine the glucose metabolic index (Rg) in **A**) lean and **B**) obese mice. The horizontal doted line in Panel B leftmost panel is the mean Rg of lean Par^f/f^ mice. The line was used to illustrate the effects of lean versus DIO mice. **C**) Gastrocnemius expression of total and phosphorylated forms of Akt were determined in basal and following acute insulin injection (1U/kg). Phosphorylated Akt isoforms were normalized to total Akt. **D**) Hexokinase II (HKII) expression was analyzed by immunoblotting in cytosolic and mitochondrial cellular fractions of the gastrocnemius muscle during basal and insulin-stimulated conditions to test the hypothesis that Parvin is required for the movement of HKII from the cytosol to the outer mitochondrial membrane. GAPDH and VDAC1 were used as indicators of cytosolic and mitochondrial fractions, respectively. **E**) muscle glycogen content. Data are presented as mean ± SE. n=4-11/group. T tests were performed in Panels A and B. Panel C and D, Two-way ANOVA with insulin and genotype as factors were run to test statistical differences between groups. *p<0.05, **p<0.01, ****p<0.0001.

### ***α***-Parvin deletion blunts aerobic exercise capacity and decreases mitochondrial respiratory flux in SkM

SkM is a major organ that contributes to whole-body energy expenditure. The observed deficits in insulin sensitivity and glucose metabolism in mParKO mice led us to question whether whole-body energy metabolism was influenced by muscle Parvin by using comprehensive mouse phenotyping coupled with indirect calorimetry (Fig. 6A). mParKO mice showed similar levels of food intake, energy expenditure, and locomotor activity compared to Par^f/f^ mice during both the light and dark cycles (Fig 6B-D). Interestingly, when given access to a running wheel to increase metabolic demand in SkM, mParKO mice voluntarily ran less total distance per day and ran at a slower speed (e.g., an estimate of exercise intensity) than Par^f/f^ mice (Fig. 6E,F). A small increase in ambulatory non-wheel activity was shown in the mParKO mice when wheels were unlocked, further suggesting that these mice spent less time on the running wheels. To determine aerobic exercise capacity, we subjected mice to an incremental exercise stress test (Fig. 6H). The mParKO mice showed reduced exercise capacity illustrated by running a shorter distance during the exercise test compared to Par^f/f^ mice (Fig 6H). This result suggests that the mParKO mice have a defect in muscle function that reduces exercise tolerance. Given these findings, we next tested whether aerobic exercise capacity was related to an intrinsic impairment in muscle oxidative function or a structural defect in muscle architecture by first performing *ex-vivo* measurements of oxygen consumption in permeabilized muscle fiber bundles. Isolated fiber bundles were supplemented with glutamate/malate to activate mitochondrial complex I, glutamate/malate plus succinate to activate mitochondrial complex II, or malate plus palmitoylcarnitine to measure fatty acid oxidation. mParKO mice showed significantly lower Complex I and Complex II activation, with oxygen consumption levels being about half of that observed in the Par^f/f^ controls (Fig. 6I). There was no difference observed in fatty acid oxidation or mitochondrial content (as indicated by citrate synthase activity) between genotypes (Fig. 6J). To test whether muscle structure was implicated in decreased muscle function and respiration, we analyzed the morphology of gastrocnemius tissue from the Par^f/f^ and mParKO mice (Fig. 6K). H&E staining revealed that mParKO mice exhibited a higher frequency of central nuclei compared to the Par^f/f^ mice (Fig. 6L), indicating muscle cells that are injured or regenerating (54). This phenotype was further corroborated using F-actin staining which revealed loss in fiber alignment in the mParKO mice and disruption of the F-actin fiber structure (Fig 6K). Thus, α-Parvin maintains muscle fiber structure and cellular organization which is required for normal mitochondrial respiration and muscular function.

**Figure 6.**
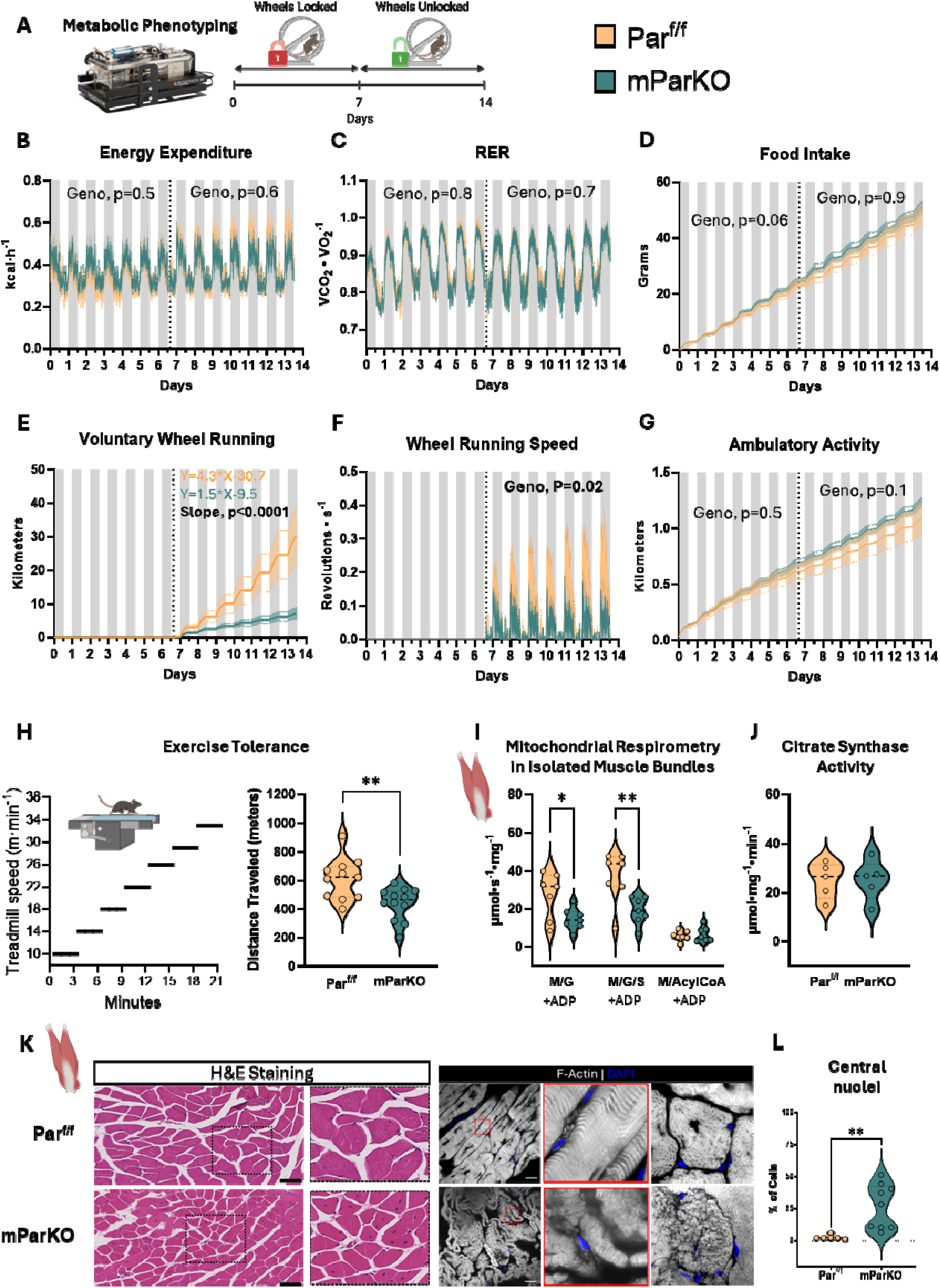
– Deletion of muscle _α_-Parvin impairs aerobic exercise tolerance and mitochondrial substrate utilization. **A**) Whole-body energy metabolism and physical activity were determined using the Promethion Core® metabolic cage system. Measurements were collected for a total of 14 days starting when mice were 16 weeks old. The first 6.5 days, running wheels were locked to prevent voluntary running activity. Thereafter the wheels were unlocked and measurements were recorded for 7 days. Metabolic and activity readouts included **B**) Energy expenditure, kcal·h^-1^; **C**) RER, respiratory exchange ratio; **D**) Cumulative food Intake; **E**) Cumulative wheel running activity; **F**) Wheel running speed; and **G**) Non-wheel ambulatory activity. **H**) Peak exercise tolerance was determined using a single lane motorized treadmill. Speed was increased every three minutes as indicated until mice could no longer maintain the running speed. The distance traveled is used as a readout of exercise tolerance. **I**) Gastrocnemius muscle bundles were isolated and incubated in substrate supplemented media to quantify muscle oxygen consumption. Malate (M) and Glutamate (G) were used to assess complex I mediated respiration; M, G, and Succinate (S) cocktail determines complex II consumption; and M plus Acyl-CoA assesses fatty acid fueled oxygen consumption. **J**) Citrate synthase activity was measured in gastrocnemius muscle as a proxy for mitochondrial content. **K**) H&E staining of gastrocnemius muscle was used to visualize muscle nuclei and gross morphological characteristics. F-actin was visualized using AF-647–phalloidin. **L**) Muscle central nuclei percentage was determined by counting of cells with and without central nuclei throughout a whole slide. Individual mice are indicated by dots on the violin plots. N=7 to 12 per genotype. A generalized linear model with body weight as a covariate was run for panel B, C, & F using CalRv2 online software (https://calrapp.org/<u>#</u>). Statistical differences for the cumulative measurements of food intake, wheel running, and ambulatory cage activity were determined using a linear regression model via GraphPad Prism software. The model was run for the wheels locked and wheels unlocked phases, separately. T tests were performed in Panels K, I, J, and L.

#### α*-Parvin deletion impairs muscle actin remodeling and GLUT4 membrane recruitment*

Since α-Parvin was required to maintain muscle fiber architecture we next asked whether the deficits in glucose uptake and metabolism was related to actin cytoskeleton dysregulation. α-Parvin mediates organization of the actin cytoskeleton by signaling through Rho-GTPases and inhibition of cofilin. Rho-GTPase activities are tightly balanced for normal actin polymerization. Thus, we measured the activity of Rac1, Cdc42, and RhoA in gastrocnemius muscle of mParKO and Par^f/f^ mice. We observed increased Rac1 and RhoA (Fig. 7A) but not Cdc42 (data not shown) activities in the mParKO mice compared to the Par^f/f^ controls. We also observed increased insulin-stimulated inhibitory phosphorylation of cofilin in mParKO compared to Par^f/f^ mice consistent with decreased actin turnover and dynamics (Fig 7B). Normal actin cytoskeleton organization is a critical requirement for muscle glucose uptake via glucose transporter 4 (GLUT4) (55). The primary upstream signal that promotes GLUT4 translocation to the plasma membrane is insulin signaling via its receptor. Therefore, we measured GLUT4 membrane localization of muscle cells in response to an acute (10 min) insulin injection (1U/kg) in 5-h fasted mice using super resolution confocal microscopy (Airy Scan 2). mParKO mice showed a reduction in GLUT4 in the membrane space of the damaged muscle fibers as defined by Caveolin3 compared to Par^f/f^ mice (Fig 7C,D). These data suggest that α-Parvin-driven actin organization and cellular structure is required for GLUT4 membrane translocation and this is likely the primary molecular mechanism underpinning muscle insulin resistance in mParKO mice.

**Figure 7.**
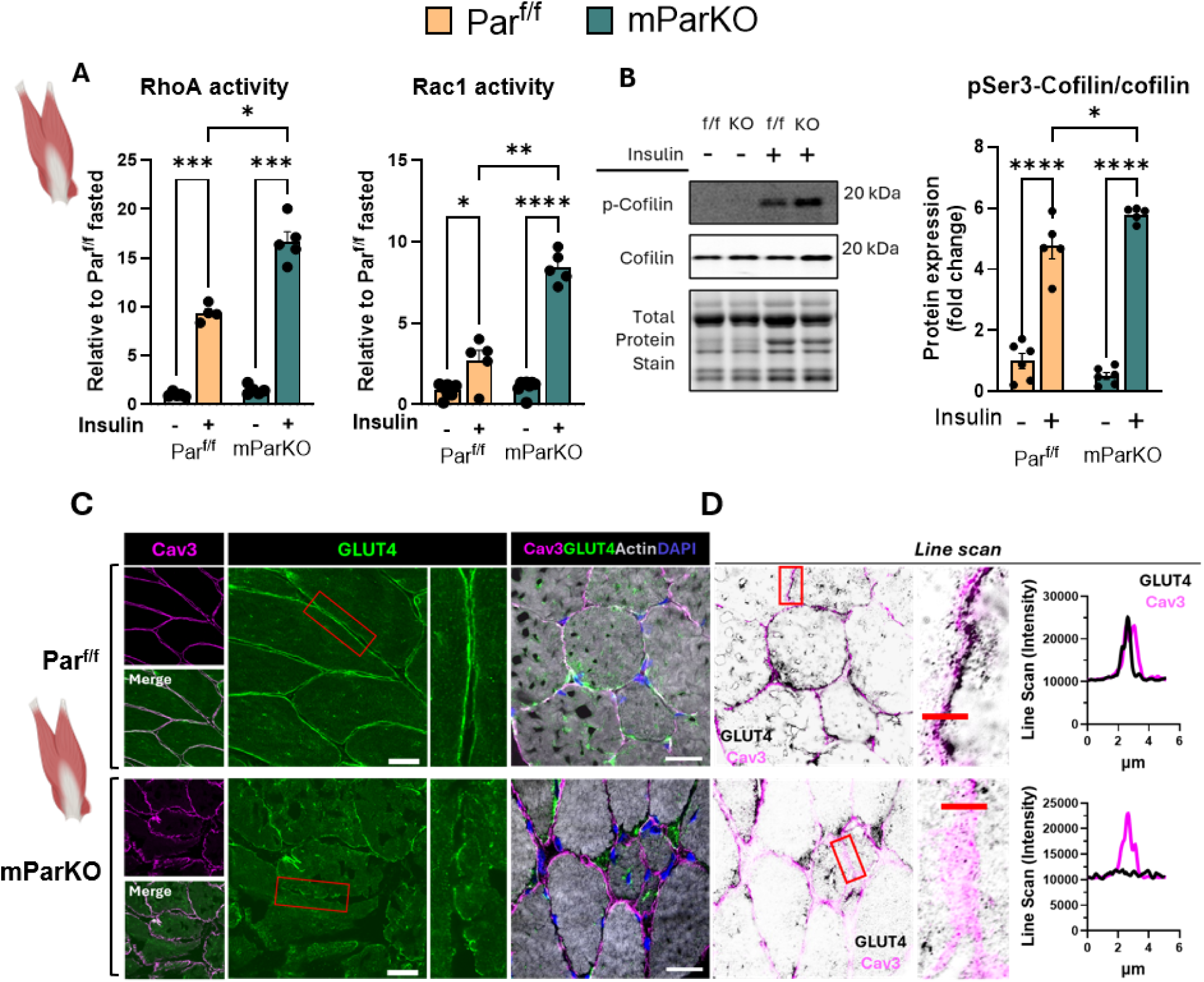
_α_-Parvin regulates skeletal muscle actin remodeling and GLUT4 membrane recruitment. **A**) RhoA and Rac1 activity assays were determined in the basal and insulin-stimulated state in gastrocnemius muscle. **B)** Immunoblotting for total and phosphorylated cofilin was determined in gastrocnemius muscle in basal and insulin-stimulated conditions. The phosphorylated cofilin densitometr was normalized to total cofilin. **C**) Sections of gastrocnemius muscle were collected from the mice indicated and fixed in a solution of 10% formalin paraformaldehyde in phosphate buffered saline or snap frozen in isopentane at liquid nitrogen temperature and affixed to cork blocks with OCT. Fixed tissue were embedded in paraffin and sectioned and frozen tissues were cryo-sectioned and mounted. Antibodies against Cav3, GLUT4, and AF-647–phalloidin (F-actin) were used to determine GLUT4 cellular localization. Confocal muscle images were collected with confocal microscopy at super-resolution using a Zeiss LSM 980 confocal microscope equipped with an inverted Axio Observer 7 and Airyscan 2 detector. **D**) Airyscan super-resolution images were acquired under identical settings for all groups and images. Acquisition and 2D Airyscan processing of acquired images was done using ZEN Blue software (Carl Zeiss). Line Scan profiles of fluorescence intensity along an annotated line was performed using the plot profile function in Fiji/ImageJ. N=4 to 6 mice per group. Panel A and B, Two-way ANOVA with genotype and insulin as factors was run to test for between group differences. *p<0.05, **p<0.01, ***p<0.001, ****p<0.0001.

## Discussion

In this study, we investigated the role of the scaffold protein α-Parvin in muscle and liver insulin resistance. We utilized conditional knockout mouse models to show that α-Parvin deletion in hepatocytes was mostly dispensable for liver insulin sensitivity in lean and obese conditions, however deletion of α-Parvin in SkM impaired muscle glucose uptake and mitochondrial respiration that was not further amplified by diet-induced obesity. Mechanistically, we find that SkM α-Parvin regulates insulin-stimulated GLUT4 membrane recruitment by controlling F-actin organization through the GTPase-cofilin pathway. These data highlight that during increased nutrient flux rates, α-Parvin-dependent actin turnover is required for muscle glucose uptake and oxidation.

The IPP complex is central to the integrin adhesome network, and it is integral to liver development and regeneration (56, 57). However, little is known about how components of the IPP regulate liver insulin resistance which is a key feature of metabolic associated steatotic liver disease. In our studies, the insulin clamp was used to determine the role of α-Parvin in hepatic insulin action in vivo. We found minimal effects of hParKO on hepatic insulin sensitivity in lean or DIO mice. Similarly, we found no differences in liver lipid content between Par^f/f^ and hParKO mice. In contrast, previous data show that hepatocyte-specific ILK knockout mice have reduced liver steatosis, and these mice have full sensitivity to the suppressive effects of insulin on glucose production (31). In the fasted state, DIO hParKO mice had elevated blood glucose, which was related to increased glycogenolysis. Moreover, insulin’s suppression of glycogenolysis was attenuated in the DIO hParKO mice. Our findings suggest that ILK and α-Parvin may act independently in the context of liver insulin action. Along these lines, prior evidence shows that liver ILK deletion accelerates liver regeneration, while PINCH deletion does not (58). Our model of DIO is a metabolic stress characterized by minimal liver injury. Whether α-Parvin KO phenocopies the loss of PINCH or ILK in more extreme stressed (e.g., injury) conditions requires additional studies.

The interactions between the ECM, integrins and integrin binding proteins have been implicated in insulin sensitivity (19, 30–32, 59–61). In this study we show that SkM glucose uptake and mitochondrial respiration require α-Parvin during high glucose flux states (insulin stimulation or exercise). These metabolic deficits in glucose uptake and metabolism originated from the muscle and contributed to reduced whole-body insulin sensitivity and exercise capacity. Our findings are in contrast to muscle specific deletion of ILK which has been shown to enhance muscle glucose uptake in obese mice (19). These diametrically opposing metabolic phenotypes of IPP complex component deletions in the same tissue highlight that we do not fully understand how Parvin and ILK regulate cellular functions. Indeed, obese SkM ILK KO mice showed enhanced insulin-stimulated Akt signaling, which was not observed in mParKO mice. In contrast, others have shown that germline deletion of ILK in skeletal muscle can lead to progression muscular dystrophy which is typically associated with insulin resistance (62). The differential activation of critical downstream signaling cascades in the absence of Parvin or ILK likely shows that in complex cells, like skeletal muscle that lack a classic focal adhesion, the IPP complex may work as a functional unit.

Another notable finding from this work is that SkM deletion of α-Parvin reduced the mitochondrial localization of HKII during insulin stimulation. HKII is a crucial step in determining glucose uptake, as it is responsible for creating the downhill concentration gradient that favors glucose transport into the cell. It irreversibly commits glucose to glucose-utilizing pathways such as glycolysis in SkM. HKII localization with the outer mitochondrial membrane may facilitate glycolytic flux directly into the TCA cycle (53). In plants, mitochondrially bound hexokinase interacts with actin and disruption of F-actin compromised glucose-dependent functions of hexokinase (63). As a corollary, we found a marked decrease in mitochondrial respiration in the presence of complex I substrates in the muscle fibers of mParKO mice. Fatty acid supplemented respiration was not reduced in mParKO mice. Fatty acids feed substrates to complexes II and III, suggesting that mitochondria of mParKO muscle may compensate for complex I deficiency by increasing electron flux through complexes II and III. Given that these mice have markedly reduced SkM glucose uptake, we hypothesize that muscle mitochondria in mParKO mice may increase their reliance on fatty acid substrates to compensate for reduced glycolytic substrates. The reduced capacity to use glucose as a fuel suggests that mParKO mice exhibit a form of metabolic inflexibility. The defect in the regulation of the actin cytoskeleton in mParKO mice is the likely contributor to the reduced capacity to accommodate the high glucose flux rates of insulin stimulation and exercise. Indeed, mParKO mice showed an increase in the phosphorylation of cofilin which inactivates its F-actin severing function, which may contribute to a rigid actin cytoskeleton. Chemical disruption of F-actin has been shown to impair mitochondrial fission (64), resulting in potentially damaged or suboptimal functioning mitochondrial. Along these lines, movement of HKII to the mitochondria was shown to be involved in mitophagy regulation, which may be influenced by RhoA kinase activity (65, 66).

Similar to the mParKO mice in this study, β-Parvin KO mice show exercise intolerance and impaired sarcomere assembly in cardiomyocytes (67). Taken together, these data show that α-Parvin not only regulates actin-mediated glucose uptake but influences mechanisms regulating glucose utilization and mitochondrial respiration.

GLUT4 in skeletal muscle is transported along tracks of microtubules which was recently shown to be disrupted in metabolic disease (68). Here, we present a model in which deletion of Parvin, a major regulator of actin cytoskeleton dynamics, results in failure of GLUT4 membrane translocation due to actin disruption as well as major metabolic defects. We find significant dysregulation of known Parvin signaling proteins like RhoA and Rac1 consistent with recent findings that Rac1 is required for normal skeletal muscle actin organization, GLUT4 membrane translocation, and that its hyperactivation in muscle is detrimental to the process of glucose uptake and storage (69, 70). The precise mechanism whereby Parvin regulates actin-driven muscle structure or GLUT4 trafficking requires further investigation. For example, it remains to be determined whether the mechanism(s) is due to a direct interaction between actin and Parvin’s actin binding domains or indirect through Parvin’s actions on RhoGTPase signaling.

Nonetheless, our findings collectively support that these molecular mechanisms in the muscle tightly control actin dynamics, which plays a previously unrecognized direct role in metabolic adaptation and glucose metabolism.

In summary, our studies show that α-Parvin is mostly dispensable for the maintenance of hepatic glucose homeostasis but is required for muscle glucose uptake during insulin stimulation and is necessary for exercise tolerance. Mechanistically, α-Parvin regulates actin cytoskeleton organization and normal muscle structure which is required for GLUT4 membrane recruitment and respiratory function. These findings demonstrate that integrin binding proteins link cell structure to metabolism via distinct tissue-specific mechanisms.

## Acknowledgments

We acknowledge the following Vanderbilt University (VU) and Vanderbilt University Medical Center (VUMC) core facilities: VUMC Hormone Assay & Analytical Services Core (NIH DK135073 and DK020593), VU Metabolic Mouse Phenotyping Center [VMMPC (NIH DK135073; www.vmmpc.org)], and Translational Pathology Shared Resource (NCI/NIH Cancer Center Support Grant 5P30 CA68485-19). We thank Tasneem Ansari, Staci Bordash, Teri Doss, Alicia Kellarakos, Carlo Malabanan for their assistance with surgical procedures and clamp studies. We also thank Merrygay James and Freya James for technical assistance.

