## Supplemental Figures for "α-Parvin Promotes Glucose Uptake and Metabolism in Skeletal Muscle with Minimal Influence on Hepatic Insulin Sensitivity"

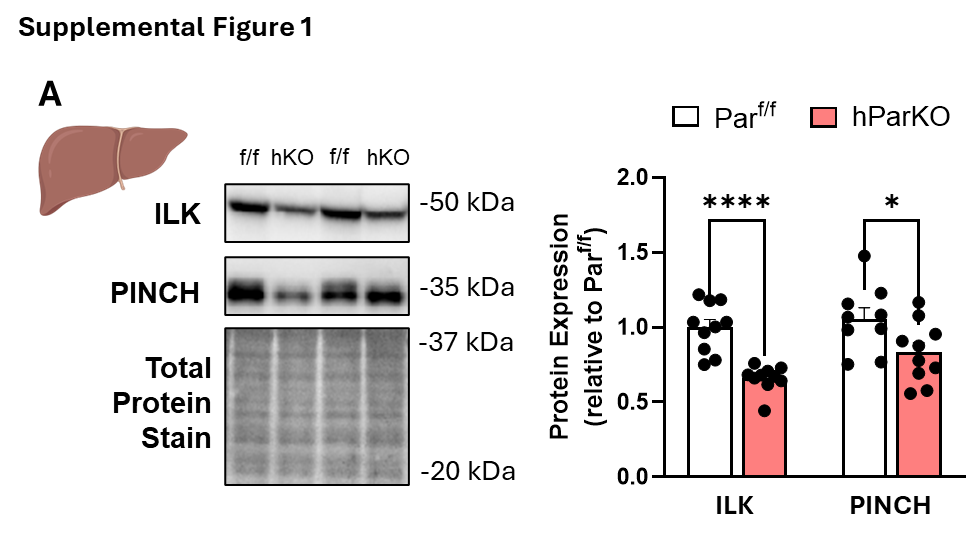

**Supplemental Figure 1** – Deletion of alpha-Parvin in hepatocytes results in a decrease in ILK and PINCH content. Immunoblotting was performed on liver lysates from mice with hepatic specific deletion of alpha-Parvin versus littermate controls. Independent samples T tests were conducted to test differences Data are presented as mean ± SE. n=9-10/genotype.

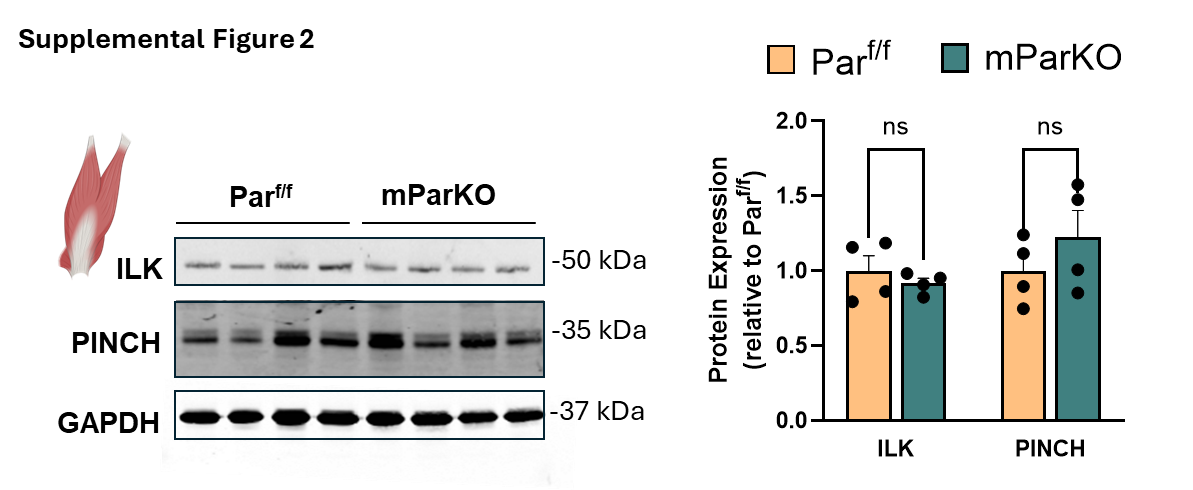

**Supplemental Figure 2** – Deletion of alpha-Parvin in skeletal muscle does not alter the protein content of ILK or PINCH. Immunoblotting was performed on gastrocnemius lysates from mice with skeletal muscle specific deletion of alpha-Parvin versus littermate controls. Independent samples T tests were conducted to test differences Data are presented as mean ± SE. n=4/genotype.
